# Evidence that Runt Acts as a Counter-Repressor of Groucho during *Drosophila melanogaster* Primary Sex Determination

**DOI:** 10.1101/648832

**Authors:** Sharvani Mahadeveraju, James W. Erickson

**Author notes:** National Institute of Diabetes and Digestive and Kidney Diseases, NIH, Bethesda, MD.

## Abstract

Runx proteins are bifunctional transcription factors that both repress and activate transcription in animal cells. Typically Runx proteins work in concert with other transcriptional regulators, including co-activators and co-repressors to mediate their biological effects. In *Drosophila melanogaster* the archetypal Runx protein, Runt, functions in numerous processes including segmentation, neurogenesis and sex determination. During primary sex determination Runt acts as one of four X-linked signal element (XSE) proteins that direct female-specific activation of the establishmen promoter (*Pe*) of the master regulatory gene *Sex-lethal (Sxl*). Successful activation of *SxlPe* requires that the XSE proteins overcome the repressive effects of maternally deposited Groucho (Gro), a potent co-repressor of the Gro/TLE family. Runx proteins, including Runt, contain a C-terminal peptide, VWRPY, known to bind to Gro/TLE proteins to mediate transcriptional repression. We show that Runt’s VWRPY co-repressor-interaction domain is needed for Runt to activate *SxlPe*. Deletion of the Gro-interaction domain eliminates Runt-ability to activate *SxlPe*, whereas replacement with a higher affinity, VWRPW, sequence promotes Runt-mediated transcription. This suggest that Runt activates *SxlPe* by antagonizing Gro function, a conclusion consist with earlier findings that Runt is needed for *Sxl* expression only in embryonic regions with high Gro activity. Surprisingly we found that Runt is not required for the initial activation activation of *SxlPe*. Instead, Runt is needed to keep *SxlPe* active during the subsequent period of high-level *Sxl* transcription suggesting that Runt helps amplfy the difference between female and male XSE signals by counterrepressing Gro in female, but not in male, embryos.

## Introduction

Cell fate decisions are commonly made in response to small quantitative differences in signal molecules. Often such signals are rendered only for brief periods during early development but lead to distinct and permanent cell fates. Sex determination in *Drosophila* is a well-defined example of a cell fate decision where a transient two-fold concentration difference in the proteins that define X-chromosome dose leads to the distinct male and female fates (reviewed in (Cline and Meyer 1996; Salz and Erickson 2010)). Four X-linked genes, *scute (sc), sisterlessA (sisA), unpaired (upd)* and *runt (run)* comprise the known X-chromosome signal elements or XSEs (Cline 1988; Duffy and Gergen 1991; Sanchez *et al.* 1994; Sefton *et al.* 2000). The XSEs function collectively to ensure that two X-chromosomes leads to the activation of the master regulatory gene *Sex-lethal (Sxl)* and thus to the female fate, whereas a single X-chromosome leaves *Sxl* inactive leading to male development (Cline 1988; Erickson and Quintero 2007). The molecular target of the XSEs is the female-specific *Sxl* establishment promoter, *SxlPe* (Keyes et al. 1992; Estes et al. 1995). In females, *SxlPe* is activated by the two-X dose of XSEs during a 30-40 minute period just prior to the onset of cellularization which occurs about 2:10-2:30 hrs after fertilization (Barbash and Cline 1995; Erickson and Quintero 2007; Lu *et al.* 2008; Li *et al.* 2011). The *Sxl* protein products produced from the brief pulse of *SxlPe* activity engage a positive autoregulatory pre-mRNA splicing loop that thereafter maintains *Sxl* protein production from the transcripts made by the constitutive *Sxl* maintenance promoter, *SxlPm* (Cline 1984; Bell *et al.* 1988; Keyes *et al.* 1992; Nagengast *et al.* 2003; Gonzalez *et al.* 2008). In male embryos, the one-X dose of XSEs is insufficient to activate *SxlPe*. Consequently the transcripts from *SxlPm* are spliced by default so as to produce nonfunctional truncated *Sxl* protein.

The four XSE elements are necessary for proper *Sxl* expression but differ in their sensitivities to gene dose and in their molecular effects on *SxlPe* (Cline 1993). The two “strong” XSEs, *sc* and *sisA*, encode transcriptional activators essential for *SxlPe* expression in all parts of the embryo (Torres and Sanchez 1991; Erickson and Cline 1993; Walker *et al.* 2000). The two “weak” XSEs *upd* and *runt* govern *SxlPe* expression in a broad region in the center of XX embryos, but neither gene is needed for expression at the embryonic poles (Duffy and Gergen 1991; Kramer *et al.* 1999; Avila and Erickson 2007). Changes in *sc* and *sisA* gene dose have dramatic effects on *Sxl* expression and consequently on viability (Cline 1988; Cline 1993). Loss of one copy of each of *sc* and *sisA* is strongly female lethal due to the failure to efficiently activate *SxlPe.* Reciprocally, simultaneous duplication of both genes is strongly male-lethal because *SxlPe* is activated in male embryos bearing an extra dose of *sc*^+^ and *sisA*^+^.

In contrast to *sc* and *sisA*, both *upd* and *runt* are relatively insensitive to changes in gene dose (Duffy and Gergen 1991; Torres and Sanchez 1992; Cline and Meyer 1996; Kramer *et al.* 1999; Sefton *et al.* 2000). Double heterozygotes between *upd* or *runt* and either of the strong XSEs show comparatively modest effects on *Sxl* expression and on female viability. Duplications of *upd*^+^ or *runt*^+^ have even smaller effects on male viability as the various combinations lead to, at most, only low-level activation of *Sxl* in XY animals. In the case of *runt*, it was only possible to detect a strong effect of *runt* dose in males, after overexpression by microinjection of *runt* mRNA into embryos (Kramer *et al.* 1999).

The *upd* gene encodes a ligand for the JAK-STAT signaling pathway and its effects on *SxlPe* are mediated via the maternally supplied transcription factor Stat92E (Harrison *et al.*1998; Jinks *et al.* 2000; Sefton *et al.* 2000). Interestingly, active Stat92E is not needed for the initial activation of *SxlPe* but is required instead to keep the promoter active during the period of maximum *SxlPe* expression (Avila and Erickson 2007). Stat92E binds to several defined DNA sites at *SxlPe* and is thought to be a conventional activator of *SxlPe* transcription that augments the functions of earlier acting XSE proteins but its actual mechanism of action is unknown (Jinks *et al.* 2000; Avila and Erickson 2007).

*runt*, encodes the archetypal member of the Runx (Runt-related transcription factor) family of proteins (Duffy and Gergen 1991; Torres and Sanchez 1992). Runx proteins are highly conserved in metazoans and act, depending on the promoter context, as either activators or repressors in a diverse array of biological processes (Walrad *et al.* 2010; ITO *et al.* 2015; Hughes and Woollard 2017). Runx proteins are defined by the Runt domain, a 128 amino acid conserved DNA binding domain that binds to the consensus binding site ‘YGYGGY’ (reviewed by (Tahirov and Bushweller 2017), and by the presence of a conserved C-terminal peptide, VWRPY, that binds to corepressors of the Groucho/TLE family (Aronson *et al.* 1997; ITO 1997; Jennings *et al.* 2006). Other conserved regions of Runx proteins mediate transcriptional activation and repression independent of the Gro-TLE family (Walrad *et al.* 2010). The *runt* gene is best known for its pair-rule function in embryonic patterning, but its initial role in the fly is as an XSE to establish female-specific expression of *Sxl* in somatic sex determination (Duffy and Gergen 1991; Kramer *et al.* 1999).

In this paper we address the mechanism by which *runt* functions to regulate *SxlPe*. We build on the experiments of Kramer et al. (Kramer *et al.* 1999) who demonstrated that Runt works directly on *Sxl* rather than through an intermediary gene. Kramer et al. (Kramer *et al.* 1999) considered three general mechanisms for how Runt might control *SxlPe*. First, Runt could act as a conventional direct activator, second; it could facilitate the binding of Sc and SisA transcription factor complexes, or third; Runt could as a “quencher” of negative regulators. Several observations focused our attention on the third possibility, that Runt activates *SxlPe* by antagonizing Groucho-mediated repression of the promoter.

Maternally supplied Groucho (Gro) is a potent co-repressor of *SxlPe* that is recruited to the promoter by DNA binding repressors of the hairy/E(spl) (Hes)-family, including Deadpan (Paroush *et al.* 1994; Fisher *et al.* 1996; Jennings *et al.* 2006; Lu *et al.* 2008). Loss of Gro leads to ectopic activity of *SxlPe* in males and premature expression in females (Lu *et al.* 2008). The first connection between Runt and Gro was the correlation between the region-specific effects of *runt* on *Sxl* and the region-specific regulation of the corepressor Gro by the Torso RTK-dependent pathway. In precellular embryos, Gro is phosphorylated directly by MAPK at the embryonic poles with phosphorylation reducing the ability of Gro to repress target genes (Cinnamon *et al.* 2008; Helman *et al.* 2011). Suggestively, the regions where Gro is phosphorylated correspond to the areas where *SxlPe* activity does not depend on *runt* (Duffy and Gergen 1991; Kramer et al. 1999). This raised the possibility that Runt is needed only in regions where Gro is highly active, a conjecture supported by early experiments showing that ubiquitous activation of Torso (which leads to ubiquitous phosphorylation of Gro (Cinnamon *et al.* 2008; Cinnamon and Paroush 2008; Helman *et al.* 2011) completely bypassed the requirement for *runt* in *Sxl* expression (Duffy and Gergen 1991). Reasoning that if Runt activates *SxlPe* by interfering with Gro, it would most likely do so via its C-terminal VWRPY peptide, we created *runt* transgenes with or without Gro-interacting motifs. We found that deletion of the WRPY sequence eliminated Runt’s ability to activate *SxlPe*, but that Runt’s transcriptional activation function was restored when the higher-affinity WRPW sequence was used. Since Runt’s ability to activate *SxlPe* depends both on the presence of a functional corepressor-interacting motif, and an intact DNA binding domain, the simplest interpretation is that Runt activates *SxlPe* by acting as a “counter-repressor” of Gro function (Pinto *et al.* 2015; Vincent *et al.* 2018). We also demonstrate that Runt is needed only after the onset of *Sxl* transcription, suggesting that *runt*, like *upd* and *Stat92E* (Avila and Erickson 2007), functions to maintain *SxlPe* in an active state. We propose a model suggesting how counter-repression by Runt could both explain Runt’s role in *Sxl* regulation and answer the paradoxical question of how a sparingly dose-sensitive XSE can play a central role in X-chromosome signal amplification.

## Materials and Methods

### Fly culture

Flies were grown at 25°C on a standard cornmeal and molasses medium. *w^1118^ run3* null flies were received from Bloomington stock center (stock number 56499). *run^3^* null mutant embryos were generated from the cross between *w f run^3^/Binsinscy* females and *run^3^/Ymal^102^(run^+^)* males. All the transgene generated were maintained with two copies in *run^3^/Binsinscy* background.

### Plasmids, Vectors and transformation

The *runt-VWRPY*^+^ 10,050 bp genomic fragment, was amplified from *w^1118^* fly genomic DNA using Expand Long Template PCR System (Roche) and cloned into pCR II-TOPO TA vector (Invitrogen). An AvrII site was introduced abutting the *runt* stop codon. The fragment ends are defined by primers: 5’-GGAAAAGTGTGTGGAAAACGGTGGA and 5’-GCAACCCAAATGTCTTGTGAAATGAA. The *runt-VWRPY*^+^ construct was modified to *runt-ΔWRPY* and *runt-WRPW* using PCR to amplification to change the C-terminal amino acids. The entire *run* coding sequences, including modifications, were introduced into the genomic clone using an AscI site located in the *runt* 5’ UTR and the introduced AvrII site and confirmed by DNA sequencing. All Runt domain mutations: Cys-127-Ser and Lys-199-Ala, Arg-80-Ala, Arg-139-Ala, Arg-142-Ala, Arg-174-Ala, Arg-177-Ala mutants were generated in pCR II-TOPO TA vector using QuikChange site directed mutagenesis kit (Agilent). The wild type and the respective modifications were confirmed by DNA sequencing. All constructs were cloned, using vector derived EcoRI sites, in the pattB transformation vector kindly provided by Johannes Bischof, Basler lab, Zurich. Transgenic injections was carried out by Genetic services Inc. MA. Constructs were inserted into fly genomic attP2 site on the third chromosome by targeted φC31 mediated specific insertion (Venken *et al.* 2006).

### In situ hybridization

Embryos were collected 0 to 3hr 30 minutes after the egg laying, Fixation of embryos and *in situ* hybridization with whole mount embryos was as described (Lu et al. 2008). Embryos are mounted in 70% glycerol in PBS for imaging. Stages of embryo were detected based on number of nuclei, shape of the nuclei and cellular furrows (Foe and Albert 1983) as outlined in Lu et al. (2008). Templates for in vitro RNA transcription was made by PCR amplification with a forward primer and a reverse primer along with T3 promoter using genomic DNA from *w1118* flies. A Digoxygenin labeled antisense RNA probe was synthesized using *in vitro* transcription kit (MAXISCRIPT T3 kit, Ambion) Probe was detected using anti-Digoxygenis antibody (Roche) that cross react with NBT-BCIP solution staining the embryos. Primers used to in vitro templates were: *Sxl* forward 5’-CCCTACGTCGACGGCATTGCAGC-3’, *Sxl* reverse 5’-TAATACGACTCACTATAGG-GAATGACCCAATGGAATCG-3’ and *runt* forward 5’-AACGACGAAAACTACTGCGGCG-3’, runt reverse 5’-AATTAACCCTCACTAAAACGGTCACCTTGATGGCTTTGC-3’.

## Results

### Runt maintains but does not initiate *SxlPe* expression

Loss of *runt* function eliminates *Sxl* protein and *SxlPe* activity, as measured by *SxlPe-lacZ* transgenes, in a broad central region in early embryos but has no apparent effect on *Sxl* at the anterior and posterior poles (Duffy and Gergen 1991; Kramer *et al.* 1999). To define precisely when and where loss of *runt* affects *SxlPe* we analyzed the effects of the *run^3^* null mutation on the production of nascent transcripts from the endogenous *Sxl* locus. Nascent transcripts from *SxlPe* were visualized as nuclear dots by *in situ* hybridization using an RNA probe derived from the *SxlPe*-specific exon E1 and downstream intron sequences. Typical results are shown in Fig. 1 with Fig. 1A highlighting nascent transcripts in magnified surface views made from the centers of the whole embryos shown in Fig. 1B. As previously reported, *SxlPe*, transcripts appear in wild-type females during nuclear cycle 12 (Erickson and Cline 1998; Avila and Erickson 2007; Erickson and Quintero 2007; Lu *et al.* 2008; Li *et al.* 2011). Initial expression was mosaic with some nuclei expressing one or both *Sxl* alleles and other nuclei neither allele. By late cycle 12 nearly all nuclei express both copies of *SxlPe* and this pattern continues, with the dots becoming more intense, through cycle 13 and the first 10-15 min of cycle 14 (Fig. 1). *SxlPe* activity decreases thereafter with the nuclear dots disappearing by mid cycle 14. Null *run^3^* mutant embryos were indistinguishable from wild-type during cycle 12, however, *SxlPe* expression began to decline in cycle 13. The decline was evident as a loss of some nuclear dots in the central portions of embryos with progressively fewer nuclei expressing *SxlPe* in later cycle 13 embryos (Fig. 1). By early cycle 14, *run^3^* null mutants displayed the expression pattern characteristic of *runt* mutants carrying *SxlPe-lacZ* fusions (Duffy and Gergen 1991; Kramer *et al.* 1999): strong expression at the poles and no expression in the broad central regions of the embryos. Our observations suggest that *runt* is not required for the initial activation of *SxlPe*, but is instead needed to keep the promoter fully active during cycles 13 and 14, but only in the central regions of the embryos. In this sense, *runt* is similar to the XSE *upd* and its associated Stat92E transcription factor, which are likewise dispensable for *SxlPe* activation but required to maintain full *SxlPe* activity after cycle 12 (Avila and Erickson 2007). The “weak” XSE elements are thus both mechanistically distinct from the “strong” XSE activators *sisA* and *scute* that are needed to activate, and presumably to maintain, *SxlPe*, activity in all portions of the embryo.

**Figure 1.**
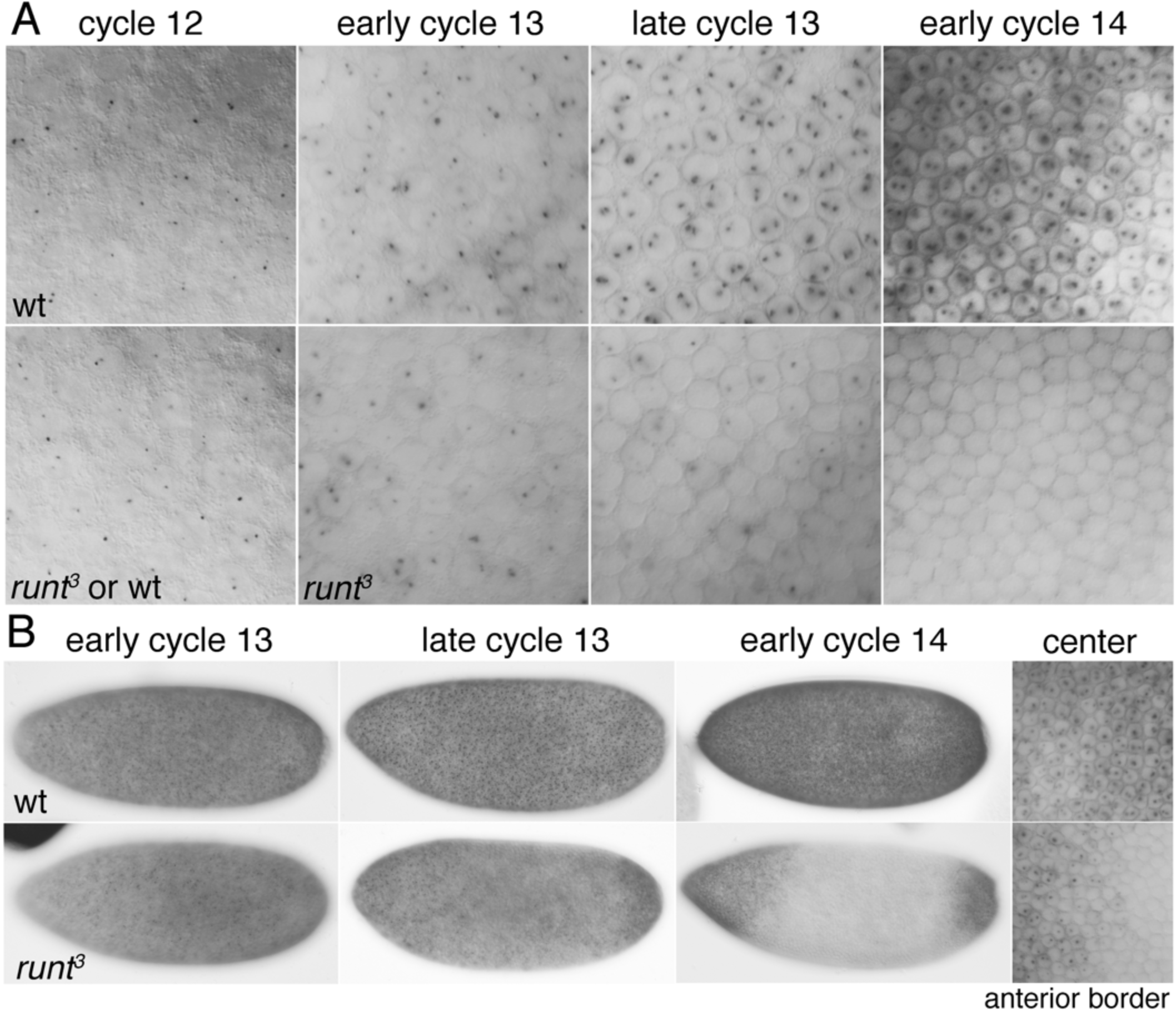
*runt* is needed to maintain but not to initiate *SxlPe* expression. Embryos were stained following *in situ* hybridization to reveal nascent and mature transcripts from *SxlPe*. Dots represent nascent transcripts from the X-linked *SxlPe*. (A) Magnified surface views from the centers of wild type (wt) and *run^3^* mutant female embryos at the indicated nuclear cycles. At cycle 12, wt and *run^3^* mutants could not be distinguished. (B) Whole embryo views of wt and *run^3^* mutant embryos at the indicated nuclear cycles. Embryos are oriented anterior to left, dorsal at top. *run^3^* null mutants displayed strong *SxlPe* expression at the poles and no expression in the broad central regions of the embryos. Right hand panels show surface views from the center of the cycle 14 wt embryo and from the anterior border between expressing and non-expressing nuclei in the cycle 14 *run^3^* embryo. Mutant embryos from the cross: *w f run^3^/Binsinscy* females X *w f run^3^/Ymal^102^ (run^+^)* males.

### Transgenes providing early *runt* function

To further analyze how *runt* regulates *SxlPe* we needed to create transgenes that express *runt* at the proper time and at appropriate levels. The *runt* gene, however, has complex regulatory regions scattered over many kilobases (Butler *et al.* 1992; Klingler *et al.* 1996) and no transgenes have yet been isolated that complement *runt* null mutations. We chose instead to isolate transgenes that reproduced the early *runt* expression pattern needed for its XSE function without concern for all of *runt’s* later functions. Using the deletion analysis of Klingler et al. (Klingler *et al.* 1996) as a guide we generated a transgene carrying a 10,050 bp genomic fragment, spanning 5,284 bp upstream of the *runt* start codon and 2,824 bp downstream of the *runt* termination codon and integrated it into the 3rd chromosome using site-specific φC31 mediated integration. We named the resulting transgene *runt·VWRPY*^+^ (Fig. 2A). We analyzed the transgenic *runt* expression pattern in the progeny of a cross between *run^3^/Binsinscy* females and *run^3^/Ymal*^102^ *(Dp run^+^)* males carrying the two copies of the *runt·VWRPY*^+^ transgene. All of embryos at or before nuclear cycle 13 expressed *runt* mRNA in patterns indistinguishable from that seen in wild type (Fig. 2A). *runt* mRNA was first detectably expressed in nuclear cycle 10. Transcripts gradually increased though cycles 13 without any visible *runt* expression in the anterior. By cycle 13 there was high-level expression in the central regions with greatly reduced mRNA staining in the posterior (Klingler and Gergen 1993). Because we could not distinguish between expression of the *runt-VWRPY*^+^ transgene and the endogenous *runt* locus in these early embryos we carefully examined 100 nuclear cycle 13 embryos, which were predicted to include ~25 homozygous *run^3^; TG/+* females, and observed no deviations from the normal *runt* expression pattern confirming that *runt-VWRPY^+^* expresses normally at this stage. While the early *runt* pattern, which is responsible for *runt’s* sex determination function (Kramer *et al.* 1999), was expressed normally from the transgenes, one quarter of cycle 14 or older embryos exhibited *runt* staining patterns that differed from the wild-type (Fig. 2B) indicating, as expected, that the transgenes lacked some regulatory sequences needed for proper expression of *runt’s* segmentation functions.

**Figure 2.**
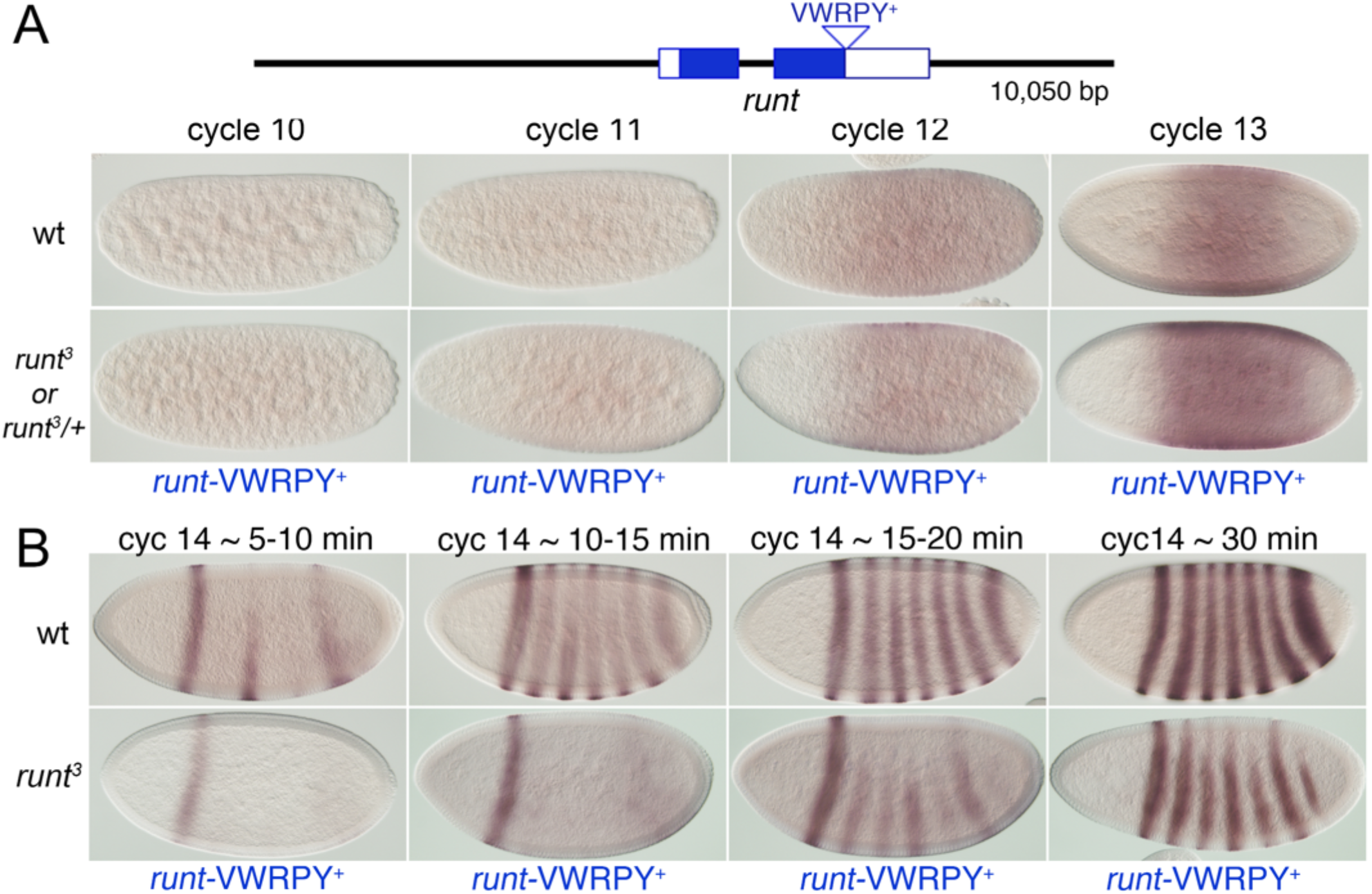
The initial *runt* expression pattern is recapitulated by the *runt-VWRPY*^+^ transgene, but not the pair-rule expression pattern. Schematic depicts of genomic DNA present in the *runt-VWRPY*^+^ transgene. Boxed regions represent coding (solid blue) and non-coding (white) sequences of the *run-RA* transcript (Flybase). The C-terminal peptide VWRPY is indicated. (A) Early *runt* expression pattern. Embryos were stained following *in situ* hybridizations to detect *runt* mRNA. Top panels show wild-type embryos at the indicated nuclear cycles. Lower panels show embryos containing one copy of *runt-VWRPY*^+^ from the cross: *w f run^3^/Binsinscy* females X *w f run^3^/Ymal^102^(run^+^); runt-VWRPY*^+^ males. Equal numbers of *run^3^* and *run^3^*/+ females and +/*Ymal^102^(run^+^)* and *run^3^/mal^102^ (run^+^); runt-VWRP*^+^ males were expected. The *run* expression patterns could not be distinguished among the embryo types. (B) *runt* pair rule expression pattern. Wild type and *run^3^* mutant embryos at the indicated times during nuclear cycle 14 stained to detect *runt* mRNA following in situ hybridization. Embryos were staged by nuclear morphology and the degree of cellularization. Stripes are located as in wild type, but are more weakly expressed, particularly in dorsal regions. Embryos are oriented anterior to the left, dorsal to the top. Genetic crosses as in (A).

### Runt-VWRPY^+^ transgenes provide XSE function

To determine if the *runt-VWRPY*^+^ transgene can provide XSE function, we asked if the transgene could restore normal *SxlPe* expression in homozygous *run^3^* mutants. We found that a single copy of the *runt-VWRPY*^+^ transgene fully complemented the *run^3^* defect as we could discern no differences in *SxlPe* activity between the *run^3^* mutant and the heterozygous female progeny of crosses between *run^3^/Binsinscy*females and *run^3^/Ymal^102^; runt-VWRPY*^+^ males (Fig. 3). Similar results were observed when the *runt-VWRPY*^+^ transgene was introduced from the female parent as expected for a zygotically acting XSE (data not shown). Taken together, the complete rescue of *SxlPe* activity in *runt* null mutants and the normal transgenic *runt* expression pattern (Fig. 2B) suggest the *runt-VWRPY*^+^ transgene produces normal or near normal levels of *runt* protein during the time when X chromosome dose is assessed.

**Figure 3.**
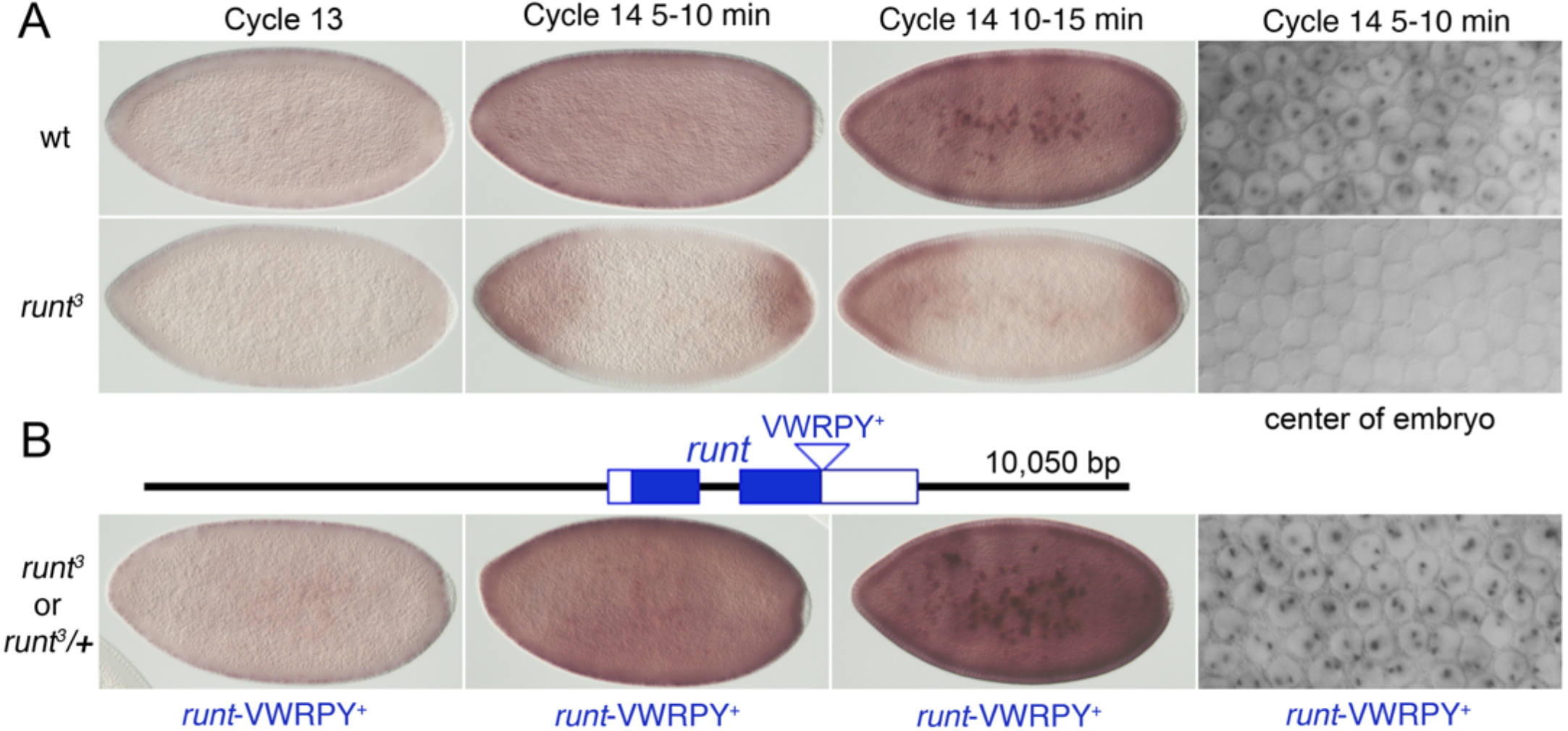
*runt-VWRPY*^+^ transgene provide full *runt* XSE function. Nascent and mature transcripts from *SxlPe* were visualized after in situ hybridization. (A) *SxlPe* expression at the indicated nuclear cycles in wt embryos and in *runt^3^* mutant females derived from the cross *w f run^3^/Binsinscy* X *w f run^3^/Ymal^102^(run^+^)*. (B) Schematic depicts the *runt-VWRPY*^+^ transgene present in single copy in the embryos shown. Since *Sxl* expression appears completely normal in *run^3^* mutants bearing *runt-VWRPY*^+^ transgenes, we could not determine if the images represent *run^3^* mutants or *run^3^*/+ heterozygotes. Cross was *w f run^3^/Binsinscy* X *w f run^3^/Ymal^102^ (run^+^); runt-VWRPY*^+^.

### DNA binding is needed for Runt to activate *SxlPe*

A requirement for Runt DNA binding in *Sxl* activation was reported by Kramer et al. (Kramer *et al.* 1999) who found that a *runt* variant carrying two amino acid changes, C127S and K199A (CK), predicted to disrupt DNA binding without greatly perturbing Runt structure, was unable to activate *Sxl* when overexpressed after microinjection of *runt* mRNA into embryos. To confirm that this finding applied to more normal levels of Runt, and to guard against the possibility that the CK amino acid replacements might otherwise alter Runt structure, we introduced the same C127S and K199A changes, as well as five single amino changes predicted to inhibit DNA binding without altering structure (Nagata and Werner 2001) into our *runt-VWRPY*^+^ transgenes (see Materials and Methods). We found that each of the amino acid changes abolished the ability of the Runt transgenes to activate *SxlPe* (data not shown) confirming that Runt’s DNA binding function is needed for its XSE function.

### Loss of Runt’s VWRPY Gro-interaction motif abolishes *SxlPe* expression

To test the significance of Runt’s Gro interactive motif in *SxlPe* activation, the WRPY portion of the motif was precisely deleted from the transgene to produce a *runt*-Δ*WRPY* derivative. (Fig. 4A). Using φC31-mediated integration, the *runt*-Δ*WRPY* transgene was inserted in the same genomic location as the wild type *runt-VWRPY*^+^ transgene. We found that Runt lacking its WRPY motif failed to rescue *SxlPe* expression in *run^3^* mutants (Fig. 4A). Indeed, the *SxlPe* pattern in *runt*-Δ*WRPY* bearing *run^3^* null mutants was indistinguishable from *run^3^* mutants alone suggesting that the Gro-interacting WRPY motif is essential for Runt to function as a transcriptional activator at *SxlPe*.

**Figure 4.**
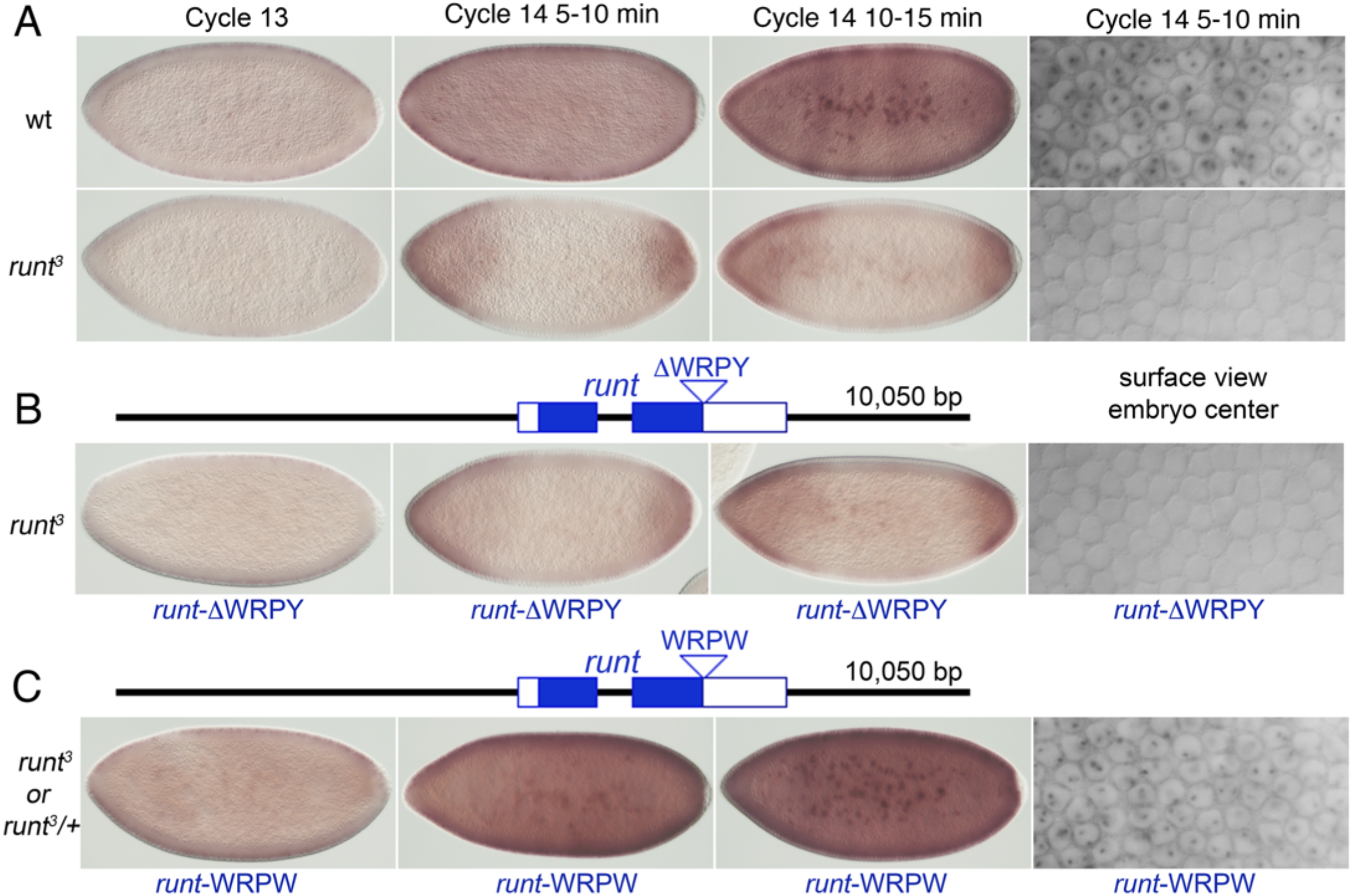
Gro-interacting C-terminal peptides are needed for Runt to activate *SxlPe*. Embryos were stained after *in situ* hybridization to reveal nascent and mature transcripts from *SxlPe*. (A) *Sxl* mRNA in wt and *run^3^* mutants at indicated nuclear cycles. Images are identical to those in Fig. 3. (B) The Gro-interacting VWRPY peptide is needed for *SxlPe* activation. Schematic shows *runt-ΔWRPY* transgene lacking the 4 C-terminal amino acids of the Gro-intracting sequence that is carried in single copy in the embryos shown. Embryos derived from the cross: *w f run^3^/Binsinscy* X *w f run^3^/Ymal^102^ (run^+^); runt-ΔWRPY*. (C) Runt protein with the high-affinity Gro binding residues, WRPW, activates *SxlPe*. Schematic shows *runt-WRPW* transgene with the Hes protein-derived WRPW Gro-interacting residues carried in single copy in the embryos shown. Embryos were from the cross: *w f run^3^/Binsinscy* X *w f run^3^/Ymal^102^ (run^+^); runt-WRPW*. Since *Sxl* expression appears normal in *run^3^* mutants bearing *runt-WRPW* transgenes, we cannot determine if the embryos shown are *run^3^* mutants or *run^3^*/+ heterozygotes.

To ensure that the failure of the *runt*-Δ*WRPY* transgene to provide sex determination reflected the loss of the WRPY motif, rather than a lack of *runt* protein, we sought a functional assay that would demonstrate the ability of the modified Runt to function in embryos in the absence of the WRPY motif. We chose to examine *fushi tarazu* (*ftz*) as previous work has shown that transcription of *ftz* is partially dependent upon *runt* activity in precelluar embryos (Tsai and Gergen 1994; Aronson *et al.* 1997; Swantek and Gergen 2004; Vanderzwan-Butler *et al.* 2007). Most important, *ftz* is activated by Runt in a partially WRPY-independent manner, as overexpressed Runt lacking the C-terminal Gro interaction domain, shows a clear activation of *ftz* expression in regions between the normal *ftz* stripes (Aronson *et al.* 1997).

We first confirmed that expression of *ftz* stripes is reduced prior to gastrulation in *run*^3^ null mutants (Fig. 5). We then showed that wild type *runt-VWRPY*^+^ transgene largely restored the endogenous *ftz* pattern. Critically, we found that the *runt*-Δ*WRPY* transgene also restored much of the normal *ftz* pattern in *run^3^* mutants, showing that the *runt*-Δ*WRPY* transgene produces functional Runt protein (Fig. 5). We note that wild type Runt was more effective at rescuing *ftz* expression than the ΔWRPY derivative. This observation, however, is entirely consistent with previous findings showing that a Runt variant lacking the C-terminal RPY residues was less effective at *ftz* activation than was the wild type when overexpressed (Aronson *et al.* 1997) as well as with the notion that *runt* likely regulates *ftz* expression by more than one mechanism (Aronson *et al.* 1997; Swantek and Gergen 2004; Vanderzwan-Butler *et al.* 2007).

**Figure 5.**
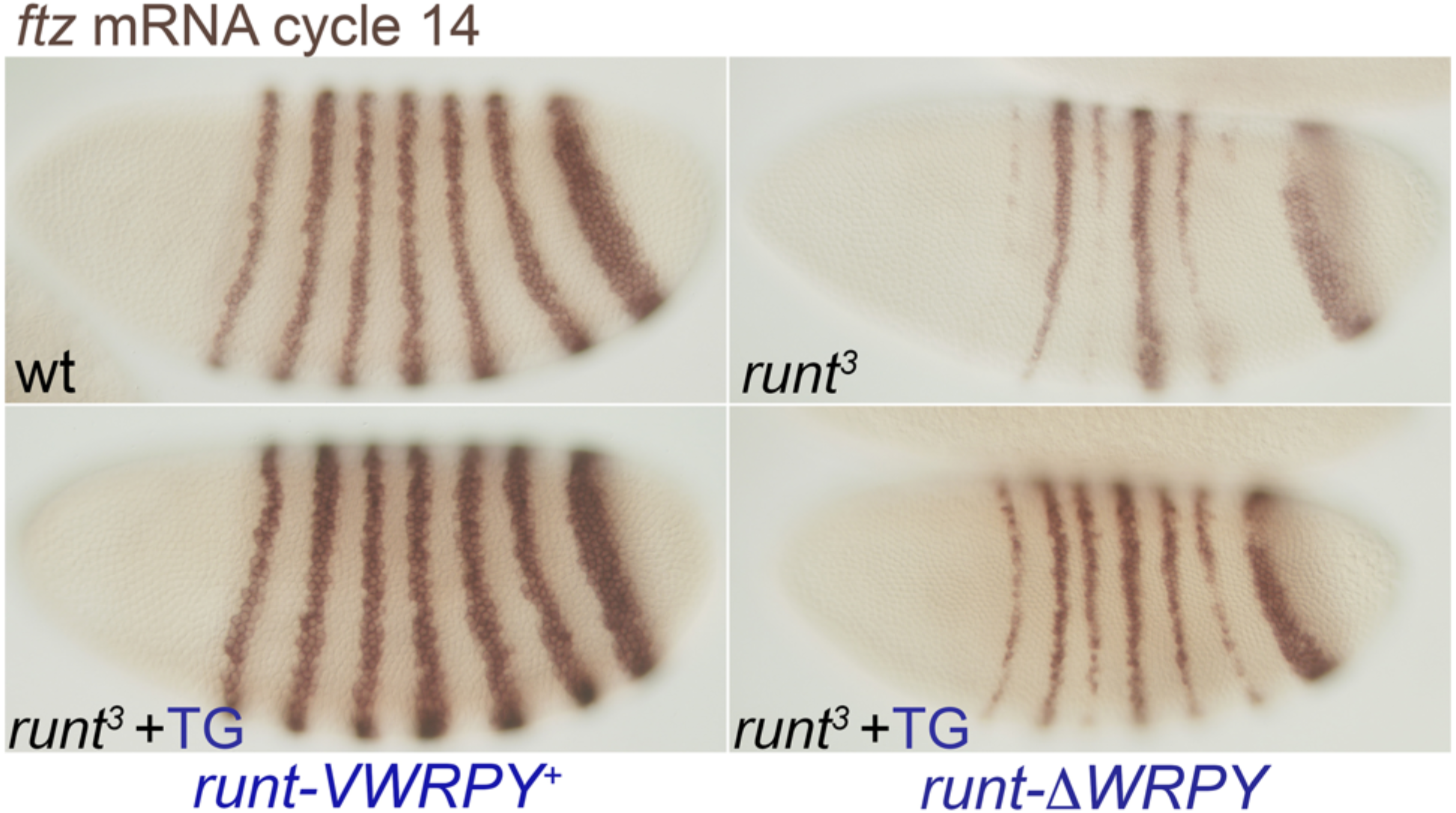
*runt-ΔWRPY* transgene retains function as it partially restores *ftz* expression in *run^3^* null mutants. Nuclear cycle 14 embryos stained following *in situ* hybridization to detect *ftz* mRNA. Top panels show *ftz* expression in wild-type and *run^3^* mutant embryos. Lower panels show *run^3^* mutants bearing one copy of *runt-VWRPY*^+^ or *runt-ΔWRPY* transgenes. Crosses were of the form: *w f run^3^/Binsinscy* X *w f run^3^/Ymal^102^ (run^+^); runt*-transgene.

### The potent Gro-interacting motif ‘WRPW’ also provides activation function at *SxlPe*

Deletion of the WRPY tetrapeptide eliminates both Runt’s interactions with Groucho (Aronson *et al.* 1997) and with its ability to activate *SxlPe* (Figure 3B). We reasoned that if Runt normally employs its VWRPY motif to antagonize Gro-mediated repression at *SxlPe* then it should be possible to substitute a different Gro interaction motif and retain Runt’s ability to activate transcription from *SxlPe*. We chose to test the well-known and potent “WRPW” Gro-interacting motif found in the dedicated repressor proteins of the *hairy-E(spl)* (HES) family. HES proteins bind Gro through their C-terminal ‘WRPW’ motif and recruit it to target gene promoters (Fisher *et al.* 1996; Fisher and Caudy 1998). The molecular interactions of Gro with WRPY and WRPW peptides are similar except that the WRPW peptide interacts with considerably higher affinity (Aronson *et al.* 1997; Jennings *et al.* 2006). We created a *runt-WRPW*^+^ transgene by changing the C-terminal ‘Y’ residue into ‘W’ and inserted the transgene into the same genomic site as the other transgenes we tested. *In situ* hybridization experiments confirmed that the *runt-WRPW*^+^ transgene restored normal *SxlPe* expression to female *run^3^* embryos (Fig. 4B). This confirms that Runt can act as transcriptional activator of *SxlPe* if its C terminus contains either a VWRPY or VWRPW co-repressor interaction motif.

## Discussion

Drosophila primary sex determination is known for its sensitivity to the concentrations of XSEs and for the rapidity of its response to the sex determination signal. During a 30-40 min period from cycle 12 through early cycle 14, *SxlPe* is turned on, its expression ramped up, and then shut down in female embryos, all while being left inactive in male embryos (Barbash and Cline 1995; Avila and Erickson 2007; Gonzalez *et al.* 2008; Lu *et al.* 2008; Li *et al.* 2011). Despite the short time available, the XSEs appear to act in at least two mechanistic stages: an initiation phase in which X dose is first sensed and a second, maintenance phase, during which the *SxlPe* activity is reinforced (Avila and Erickson 2007). The highly dose-sensitive “strong” XSE proteins, Sc and SisA, appear to act in both stages as complete loss of either, or a two-fold reduction in both, effectively eliminate *SxlPe* activity and the temperature-sensitive period for *sc* extends into cellularization (Erickson and Cline 1993; Walker *et al.* 2000; Wrischnik *et al.* 2003). Remarkably the two more weakly dose-sensitive XSE proteins, Runt and Upd, act at the second stage as both are dispensable for the initial activation of *SxlPe* but are critical for maintaining full promoter activity during cycles 13 and 14 (Fig. 1, (Avila and Erickson 2007)). A two-step model offers a possible explanation for the paradoxical notion that two critical players in this textbook example of a dose-sensitive genetic switch are themselves relatively dose-insensitive (Duffy and Gergen 1991; Torres and Sanchez 1992; Cline 1993; Sanchez *et al.* 1994; Kramer *et al.* 1999; Sefton *et al.* 2000). The exact gene dose of the weak XSE elements would not matter to male embryos if Runt and Upd, or the Stat92E transcription factor it activates, are only capable of enhancing transcription from an already active *SxlPe.* This could be the case if Runt or Stat92E are unable to bind to or function at *SxlPe* unless the promoter has already been activated by the strong XSE proteins. We note that male-specific viability is unaffected even with a total of four copies of wild-type *runt*, (one each on the *X* and *Ymal^102^* chromosomes, and two transgenic copies, unpublished data), a finding in stark contrast to what was seen with *sc* or *sisA* which are strongly male-lethal if either one is present in three copies (Erickson and Cline 1991; Erickson and Cline 1993; Cline and Meyer 1996; Wrischnik *et al.* 2003). In females, Runt plays a critical role in maintaining *SxlPe* in the on state during nuclear cycles 13 and 14; however, females would be relatively insensitive to *runt* and *upd* dose if a single copy of each gene provided enough Runt or active Stat92E to effectively reinforce the actions of Sc and SisA. In contrast, if *SxlPe* activity were partially compromised by reductions in *sc* or *sisA* dose an additional reduction in *runt* dose might exacerbate the *Sxl* expression defect leading to the observed female-lethal effects (Duffy and Gergen 1991; Torres and Sanchez 1992).

Evaluating the validity of models of dose-sensitivity requires that the molecular functions of the XSEs be elucidated. The XSE protein Sc and its maternally supplied partner, Daughterless, are bHLH transcriptional activators that bind as heterodimers to six or more sites at *SxlPe* known to important for transcription (Yang *et al.* 2001). SisA remains an enigma but appears to be a non-canonical bZIP transcription factor (Erickson and Cline 1993; Fassler *et al.* 2002). The *upd* protein signals activation of Stat92E, a maternal transcription factor that binds sequences needed for full *SxlPe* activity (Jinks *et al.* 2000; Avila and Erickson 2007; Cline *et al.* 2010). Stat proteins, like Runx proteins, tend to be relatively weak activators that require interactions with other proteins to activate transcription (Horvath 2000; Goenka and Kaplan 2011). Intriguingly, Stat92E, has been shown to function as a positive regulator of the *crumbs* enhancer, *crb518*, via a counter-repression mechanism (Pinto *et al.* 2015), raising the possibility that Stat92E could function at *SxlPe* in a manner conceptually similar to what we propose here for Runt.

Runt is a bifunctional transcription factor that activates or represses a variety of cellular targets. A common mechanism of repression involves Runt’s C-terminal pentapeptide, VWRPY, which is needed to recruit the potent co-repressor Gro to targets including *even-skipped, hairy*, and *engrailed* (Aronson *et al.* 1997; Walrad *et al.* 2010). Still other targets of Runt and Runx proteins are repressed via Gro- and VWRPY-independent mechanisms (Walrad *et al.* 2010; Walrad *et al.* 2011; Hang and Gergen 2017). Activation by Runt is best understood at *sloppy-paired-1* (*slp1*) where Runt interacts with the transcription factor Opa to bind the *slp1* DESE enhancer to drive expression in odd numbered *slp1* stripes (Swantek and Gergen 2004; Walrad *et al.* 2010; Walrad *et al.* 2011; Hang and Gergen 2017). Interestingly, deletion of Runt’s C-terminal 25 amino acids, including the VWRPY motif, prevents Runt from activating *slp1;* however amino acids other that the VWRPY motif appear to be involved as Gro appears to play no role in regulating the DESE enhancer. Here we provide evidence that Runt activates *SxlPe* by interfering with Groucho mediated repression suggesting that Runt’s role at *SxlPe* is as a counter-repressor (Pinto *et al.* 2015; Vincent *et al.* 2018). First, we showed that deletion of just the Gro-interacting WRPY sequence rendered a *runt* transgene that normally provides full XSE function, unable to activate *SxlPe* (Fig. 3A). Second, we found that a *runt* derivative containing the higher affinity Gro-interaction motif WRPW sequence from Hes-class repressors also functions as an activator of *SxlPe* (Fig. 3B). Critical to our analysis, was the finding that the *runt*-Δ*WRPY* transgene that failed to activate *SxlPe* was capable of partially rescuing the *runt*-dependent loss of *ftz* stripes (Fig. 4), a function known to be partially dependent on Runt’s WRPY motif (Aronson *et al.* 1997). We attempted to obtain additional evidence for the presence of the Runt-ΔWRPY protein in embryos using whole mount immunostaining but were unable to obtain antibody preparations that could detect wild type Runt protein. We acknowledge this limitation of our experiments, but note that deletion of a short C-terminal sequence that included the VWRPY motif did not destabilize Runt when overexpressed in Drosophila salivary glands or early embryos (Walrad *et al.* 2010). Similarly, loss of the VWRPY peptide does not destabilize mammalian Runx1 or Runx3 VWRPY mutants in cultured cells or live animals (Nishimura *et al.* 2004; Yarmus *et al.* 2006; SEO *et al.* 2012).

Our finding that Runt requires its co-repressor interaction domain to function as an activator of *SxlPe* may appear surprising; however, it is not a novel idea, having been first proposed in the paper that showed the physical interactions between Runt and Gro (Aronson *et al.* 1997) and discussed further by Kramer et al. (Kramer *et al.* 1999) and McLarren et al. (McLarren *et al.* 2000; McLarren *et al.* 2001) who proposed that Runt might interfere with Gro function at *SxlPe* in females and then actively promote further transcription. Importantly, it fits well with both the central role of Gro-mediated repression in *SxlPe* regulation (Paroush *et al.* 1994; Lu *et al.* 2008) and with a variety of published data on Gro and Runt function.

Maternally supplied Gro is recruited to *SxlPe* by DNA binding proteins including the Hes protein, Dpn. Dpn binds to three sites within 160 bp of the start of *SxlPe* transcription (Lu *et al.* 2008). While Gro is often considered a long-range repressor recent analyses has revealed that short-range repression, with Gro-binding near the promoter, as occurs at *SxlPe*, is more common (Kaul *et al.* 2014; Kaul *et al.* 2015). Loss of maternal Gro has several effects on *SxlPe.* It causes ectopic expression in male embryos and premature *SxlPe* activity in females. This suggests maternal Gro defines the initial threshold XSE concentrations needed to activate *SxlPe* and that it actively keeps the promoter off in males. In the absence of Gro, *SxlPe* appears to be expressed in direct proportion to X chromosome dose suggesting that Gro plays a central role in X-signal amplification (Lu *et al.* 2008). Antagonism of Gro function is thus a plausible means by which an XSE might regulate the *SxlPe* switch. The most suggestive prior indication that Runt might work by inhibiting Gro function was that Runt is needed for *Sxl* expression only in the broad central domain of the embryo where Gro-mediated repression is most effective. Runt is not required at the embryonic poles where Torso-signaling leads to the down regulation of Gro activity via phosphorylation (Cinnamon *et al.* 2008; Kaul *et al.* 2015). In this context, the then mysterious observation by Duffy and Gergen (Duffy and Gergen 1991), that a *torso* gain-of-function allele completely bypasses the need for *runt* in *Sxl* activation is easily explained. Expression of constitutively active torso leads to uniform phosphorylation and inactivation of Gro (Cinnamon *et al.* 2008; Cinnamon and Paroush 2008; Helman *et al.* 2011). Absent active Gro, there is nothing for Runt to counter-repress at *SxlPe*.

How might Runt inhibit Gro function? Based on our findings and those of Kramer et al. (Kramer *et al.* 1999) it would appear that Runt must bind to DNA to activate *SxlPe* suggesting that Runt likely inhibits Gro at the promoter. This would rule out a titration scheme in which Runt binds Gro and prevents it from being recruited to *SxlPe* by DNA binding repressors. Plausible mechanisms of Gro inhibition could involve local phosphorylation of Gro at *SxlPe* if Runt could recruit a protein kinase to the promoter, or direct competition with the Hes-repressors, such as Dpn, for Gro-binding (McLarren *et al.* 2000; McLarren *et al.* 2001). It is also possible that changes in Gro structure induced by Runt binding to it at *SxlPe* might inactivate Gro. An intriguing possibility is that Runt’s interaction with Gro at *SxlPe* could be mediated by an XSE or an XSE-dependent co-factor. The ability of the Drosophila Runx protein, Lozenge, to stably associate with Gro in eye development depends on its interactions with the transcription factor Cut (Canon and Banerjee 2003). While the interaction with Cut regulates Lozenge’s function as a repressor, a similar mechanism could promote a counter-repressing interaction with Gro.

A remaining mystery is where Runt binds at *SxlPe* as no specific Runt DNA binding sites have been identified near the promoter. Kramer et al. (Kramer *et al.* 1999) reported that Runt, and its CBF-β DNA binding partner, Brother (Bro), bound several 200-300 bp DNA fragments from the *SxlPe* region; however, binding specificity was tested only by competitive challenge with high-affinity consensus DNA binding sequences. Our laboratory also found that Runt, in combination with Bro (or the other CBF-β protein, Big-brother) bound a variety of *SxlPe* fragments, but we observed that binding was efficiently competed in every case by low concentrations of non-specific (poly dI-dC) competitor (unpublished data). Given the absence of obvious matches to the Runt binding site consensus at *SxlPe* and the inability to identify specific in vitro binding sites, it suggests that Runt may bind to *SxlPe* only in combination with other protein complexes.

The notion that Runt might target Gro function only after *SxlPe* has been activated offers a possible explanation for how the sparingly dose-sensitive, *runt* protein could play an important role in amplifying the two-fold difference in male and female XSE doses into a reliable developmental signal. We previously proposed a model in which female-specific dampening of Gro-mediated repression was a central part of X-chromosome signal amplification (Lu *et al.* 2008; Salz and Erickson 2010). Our focus in the earlier paper was a hypothetical feedback mechanism by which active transcription of *Sxl* reduced Gro-mediated repression of *SxlPe*. In the modified version of the model (Fig. 6), Runt, and potentially Stat92E, counteract Gro-mediated repression in female, but not in male, embryos. The central tenets of the model are that the 2X dose of the strong XSEs provides sufficient Sc and SisA to cross the threshold for *SxlPe* activation during cycle 12, but that their combined concentrations are insufficient to keep the promoter active in the face of increasing repression as the zygotic repressor Dpn accumulates and translation of maternal Gro mRNA continues. The “weak” XSEs function to counteract repression after *Sxl* transcription begins, either by further enhancing *SxlPe*, as may be the case if Stat92E functions as an activator, or by directly inhibiting Gro function by counter-repression as we propose for Runt. Signal amplification would occur because the increasing XSE protein concentrations in 2X embryos maintains the promoter in an active state, whereas the 1X dose of XSEs can never overcome the ever-increasing repression in males.

**Figure 6.**
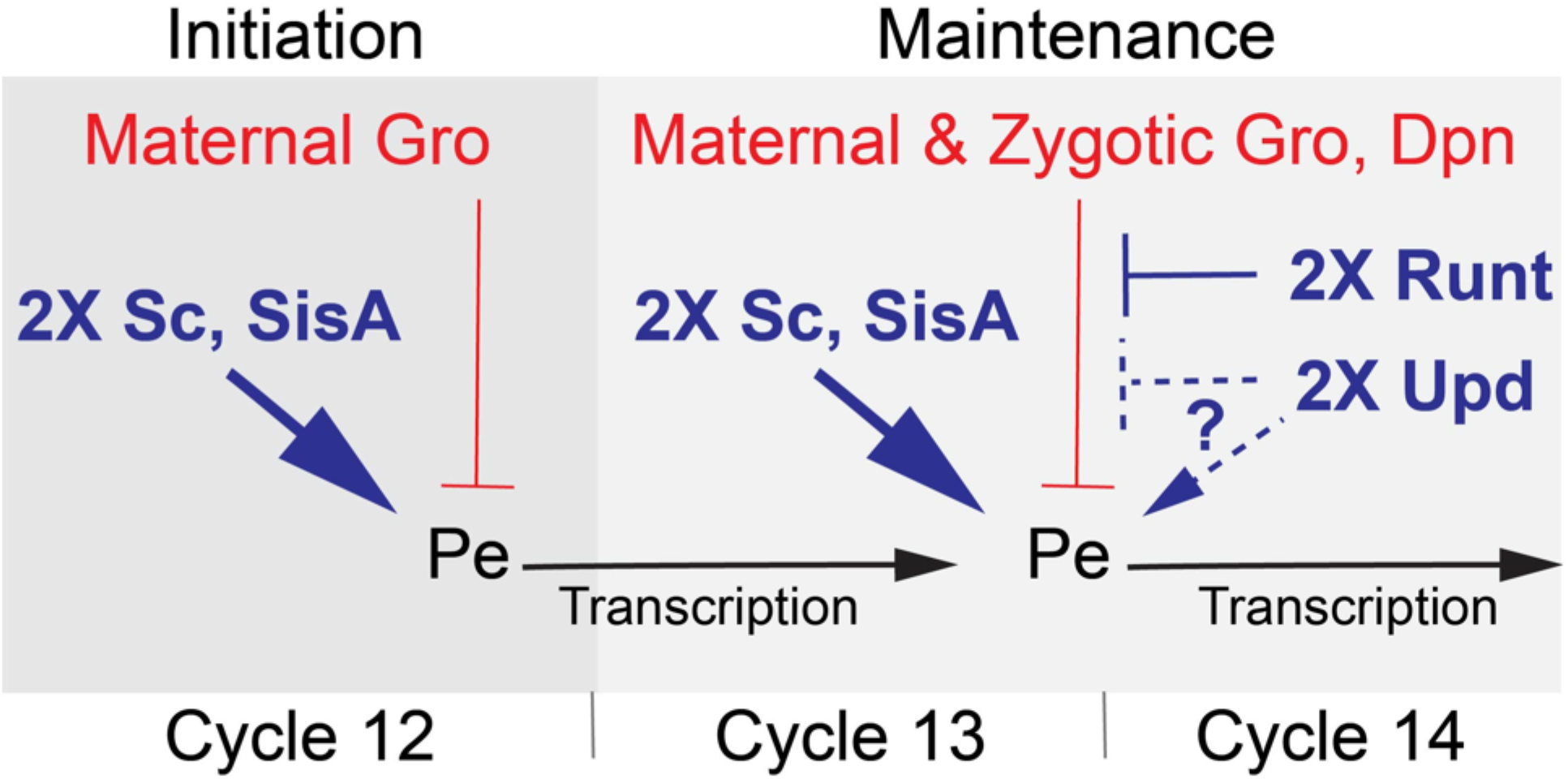
Model for regulation of *SxlPe.* In female embryos the two X dose of the XSE transcription factors, Sc and SisA, overcomes maternal Gro repression initiating expression from *SxlPe* in nuclear cycle 12. During cycles 13 and 14 increased levels of Sc and SisA, assisted by Runt and Upd, maintain *SxlPe* transcription. Runt counter-represses Gro function via its VWRPY domain. Upd, acting through the STAT92E transcription factor, may activate *SxlPe* directly or counteract repression. In male embryos, the single X doses of Sc and SisA fail to overcome Gro-mediated repression and do not activate *SxlPe*. Without *SxlPe* activation, Runt and Upd/Stat92E do not function at *SxlPe*.

Might the kind of counter-repression mechanism we propose for Runt at *Sxl* exist for other genes? Interestingly, McLarren et al. (McLarren *et al.* 2001) observed that mammalian Runx2 inhibited the ability of Hes1 and the mammalian Gro protein, TLE1, to repress an artificial promoter in cultured rat osteosarcoma cells. While the authors did not test if the Runx2 VWRPY residues were needed for relief of TLE1-mediated inhibition, they did note the apparent commonalities with Drosophila sex determination. Further analysis of genes co-regulated by Runx, Hes, and Gro/TLE family proteins should reveal whether it is common for Runx proteins to activate genes by interfering with repression.

## Acknowledgments

Stocks obtained from the Bloomington Drosophila Stock Center (National Institutes of Health P40OD018537) were used in this study. The work was supported by a grant from the National Science Foundation, #1052310, and by funds provided by Texas A&M University and by JWE.

